# Hydrogen stable isotope probing of lipids demonstrates slow rates of microbial growth in soil

**DOI:** 10.1101/2022.07.11.499392

**Authors:** Tristan A. Caro, Jamie McFarlin, Sierra Jech, Noah Fierer, Sebastian Kopf

## Abstract

The rate at which microorganisms grow and reproduce is fundamental to our understanding of microbial physiology and ecology. While soil microbiologists routinely quantify soil microbial biomass levels and the growth rates of individual taxa in culture, there is a limited understanding of how quickly microbes actually grow in soil. For this work, we posed the simple question: what are the growth rates of soil microorganisms? In this study, we measure these rates in three distinct soil environments using hydrogen stable isotope probing of lipids with ^2^H-enriched water. This technique provides a taxa-agnostic quantification of *in situ* microbial growth from the degree of ^2^H enrichment of intact polar lipid compounds ascribed to bacteria and fungi. We find that average apparent generation times in soil are quite slow (20 to 64 days) but also highly variable at the compound-specific level (6 to 1137 days), suggesting differential growth rates between community subsets. We observe that low-biomass communities can exhibit more rapid growth rates than high-biomass communities, highlighting that biomass quantity alone does not predict microbial productivity in soil. Furthermore, within a given soil, the rates at which specific lipids are being synthesized do not relate to their quantity, suggesting a general decoupling of microbial abundance and growth in soil microbiomes. More generally, we demonstrate the utility of lipid stable isotope probing for measuring microbial growth rates in soil and highlight the importance of measuring growth rates to complement more standard analyses of soil microbial communities.

**Significance:** Generation times, how quickly organisms grow and reproduce, are a key feature of biology. However, there are few measurements of microbial generation times in soil, despite the crucial importance of soil microbes to terrestrial ecosystems. By measuring the rate at which isotopically labeled water is incorporated into microbial membranes, we find that the generation times of soil microorganisms are far longer than those typically observed in culture. Surprisingly, we observe that lower-biomass soils exhibited faster growth rates than high-biomass soils. More abundant microorganisms are not necessarily the fastest growing and most soil microorganisms are slow growers. Our results underscore the importance of considering slow and variable growth rates when studying microbial communities and their contributions to ecosystem processes.

## Introduction

The rate of microbial growth is a parameter commonly invoked in biogeochemical models of carbon flux, nutrient uptake, ecosystem productivity, and other soil assessments. However, growth states of microorganisms are extremely variable, with doubling times ranging from minutes in laboratory-based culture to many months and years for organisms living in the Earth’s subsurface (1–3). Drastically different lifestyles and strategies characterize the microbial world, with oligotrophic systems sustaining slow-growing organisms on timescales far beyond those typically observed in more resource-rich environments that select for rapid growth and short generation times (4, 5).

Numerous studies have measured soil microbial biomass with the assumption that the size of the standing pool of microbial biomass is an important metric of soil productivity (6–11). However, static biomass assessments are not necessarily a measure of the influence that microorganisms might have on ecosystem processes in real time, including nutrient and carbon cycling. For instance, an environment might consist of a largely static biomass pool with low turnover, or conversely, a small biomass pool with high turnover. Measurements of total soil microbial biomass may therefore yield an incomplete picture of microbial contributions to ecosystem processes. Indeed, in plant communities, ecosystems with high standing stocks of plant biomass can often be less productive than low-biomass communities (e.g., boreal forests versus grasslands). We cannot assume *a priori* that microbial communities with larger standing biomass are necessarily more productive. Just as net primary production (NPP), not standing biomass, is used as a growth parameter in plant systems, a similar metric is required to assess the productivity of soil microbial communities.

Growth rate is a critical parameter with which to assess ecosystem function, but there remains a lack of consensus around methods for measuring *in situ* microbial growth in soil systems. As a result, microbial growth rates in soil are poorly constrained, with previously reported community-level generation times varying by several orders of magnitude (12–20). For a thorough overview of the pre-existing methods for measuring soil microbial growth rates, see Rousk & Bååth (13).

Stable isotope tracers (e.g., ^13^C, ^2^H, ^15^N) have been used to assess microbial activity in a wide variety of systems, from clinical samples to the deep biosphere (3, 21–25). Stable isotope probing (SIP) relies on the addition of an isotopically labeled tracer followed by time-series measurements to calculate the rate of tracer incorporation into biomolecules by active microorganisms. SIP, unlike methods that rely on cell counting or biomass estimation, can provide information on biosynthetic or metabolic turnover independent of both the size of the standing biomass pool and the population dynamics of the community (i.e., population growth, steady state, or decline). Thus, SIP approaches are well-suited to assess growth rates in soils, where standing biomass pools may vary widely, and population dynamics may be spatially or temporally variable.

Here, we employ a dilute deuterated heavy water (^2^H_2_O) tracer (5000 ppm ^2^H) because all organisms incorporate water-derived H into their lipids during growth, and the addition of labeled water should not select for or against the growth of any organisms, thus rendering the tracer taxa-agnostic and nutritionally neutral. The incorporation of ^2^H from ^2^H_2_O into microbial membrane lipids can be measured by gas chromatography/pyrolysis/isotope ratio mass spectrometry (GC/P/IRMS) along a time series and converted into a growth rate, as previously demonstrated for clinical samples and marine sediments (22, 23, 26–29). This method, which we abbreviate as lipidomic hydrogen stable isotope probing (LH-SIP) offers several advantages for the estimation of microbial growth rates *in situ*. First, it allows us to use reasonably short incubation times (days) to measure the generation times of microorganisms, even if they are very long, as this method only requires a fraction of an organism’s generation time to elapse to yield quantifiable growth rates. Moreover, all microorganisms routinely synthesize lipids regardless of their stage in the cell cycle or metabolic activity. Unlike nucleic acids and proteins, which can be re-synthesized for cellular maintenance and repair even in the absence of growth, lipids are less likely to require repair and thus provide a more specific measurement of membrane and cell growth (30). Isotope ratio mass spectrometers have a wide dynamic range that is well-suited for capturing the trace isotopic incorporation that is expected from exceptionally slow turnover rates. The accuracy and range of IRMS instruments in ^2^ H space allow for accurate quantification of incorporation corresponding to ∼1/1000^th^ of an organism’s generation time in the presence of as little as 1% ^2^H_2_O. Thus, LH-SIP is well-suited for the study of slow-growing microorganisms in soil.

For this study, we focused on three soils that represent different site and edaphic characteristics (Supplementary Data 1), with these soils harboring distinct microbial communities (Supplementary Figure 3). We directly measured growth rates in these soils with compound-specific LH-SIP, highlighting the utility of our approach, the relevance of measuring microbial growth rates in soil, and the broader implications of our results for understanding soil microbial dynamics.

## Results and Discussion

### Measuring the generation times of soil microorganisms

We measured ^2^H incorporation into soil microbial phospholipid fatty acid (PLFA) components at three time points during a 7-day incubation period in the presence of a dilute heavy water (^2^H_2_O) tracer. We detail these experiments in *Materials and Methods*. In brief, we incubated three soils (a sub-alpine conifer forest, prairie grassland, and an alpine tundra soil) in the presence of 5000 ppm (*δ*^2^H_VSMOW_≈ 31,000 ‰)^2^H_2_O, extracted intact polar phospholipids, and measured their fatty acid abundances by gas chromatography flame ionization detection (GC-FID) and isotopic compositions by gas chromatography isotope ratio mass spectrometry (GC-IRMS). We report isotopic values as fractional abundance (^*2*^*F*) in units of ppm where^2^F =^2^H / (^2^H +^1^H). We observed^2^H incorporation into fatty acids to range from 10^1^to 10^3^ppm in the presence of 5000 ppm^2^H_2_O tracer (Figure 1A). We inferred the turnover rate and apparent generation times of microbial lipids using a previously derived relationship (22) (Supplementary Text). In short, microbial growth (µ) is a logarithmic function of incubation time and the fractional hydrogen (^*2*^*F*) isotopic enrichment of new biomass relative to that of biomass at the start of the incubation. We report both lipid-specific turnover rates as well as the abundance-weighted mean (i.e., community-level mean) turnover rate for each soil. While the majority of the PLFAs detected are sourced from prokaryotic organisms (31), PLFAs attributable to fungi (31, 32) are also reported.

**Figure 1.**
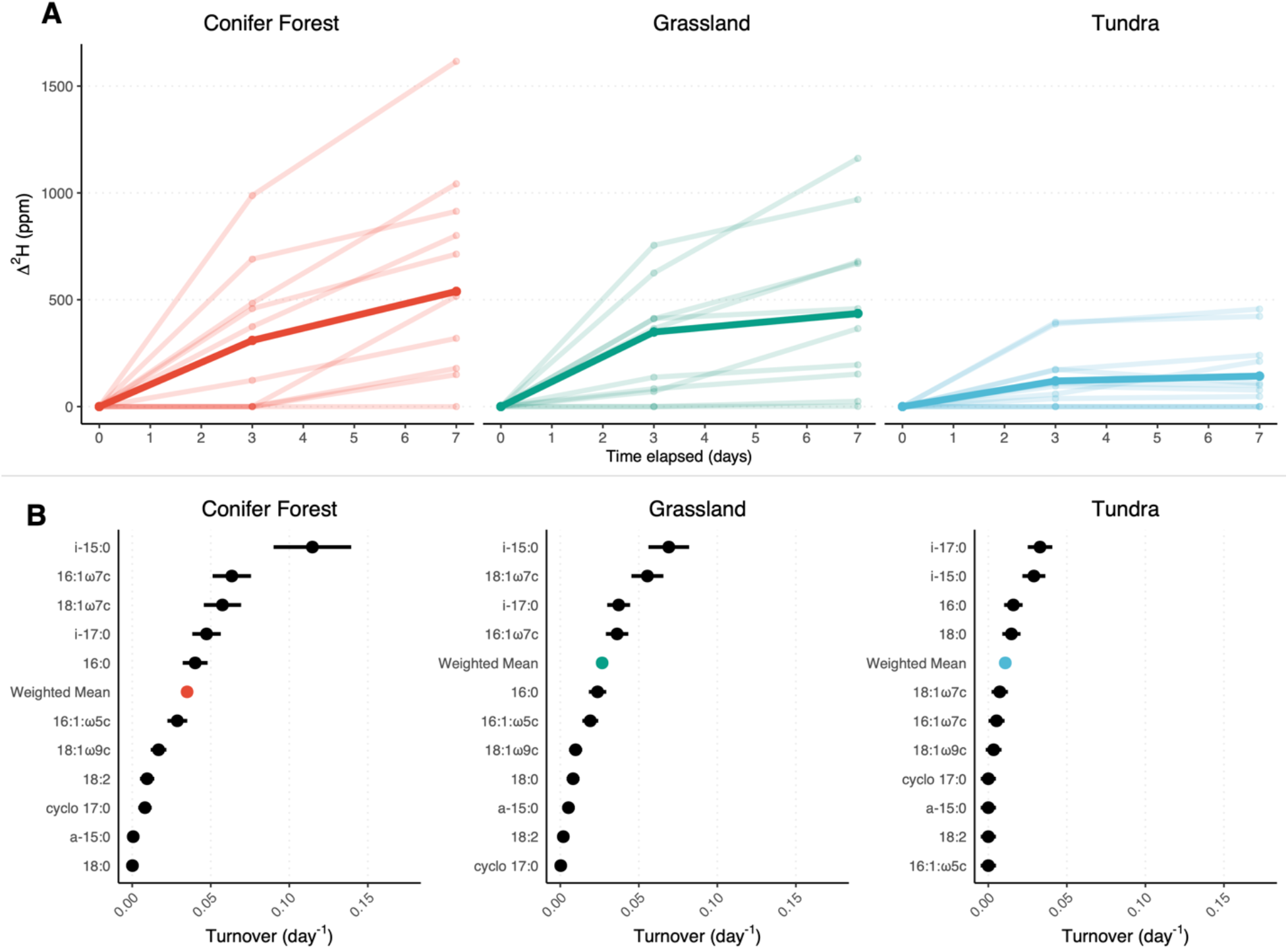
Δ^2^H enrichment (defined as change in ppm relative to the start of the incubation) of microbial biomass increases over the course of the incubation, allowing the calculation of compound-specific and weighted-mean growth rates (day^-1^) (panel A). The mean^2^H enrichment (A), weighted by compound abundance, is shown in bold. For Panel B, turnover rates were calculated using the integrated Δ^2^H between zero and seven days. Error bars correspond to the propagated uncertainty of isotope measurements and tracer assimilation (Supplementary Text). Data used to generate this figure is available in Supplementary Dataset S4.

### Rates of microbial growth in soil are slow

The grassland and conifer forest soils both exhibited slow microbial growth rates (weighted mean generation times of 26.0 d and 19.8 d respectively). Growth was far slower in the alpine tundra (weighted mean generation time of 64.2 d). The alpine tundra soil experiences the lowest mean annual temperature (MAT) of all the sites studied here, with a MAT of -3°C (33), and is characterized by short (30 – 90 day) and cool growth seasons, where soil respiration begins immediately after soils begin to thaw underneath seasonal snowpack (34). The generation times we observed under conditions analogous to the warm season (20°C) suggest that, even in the warm season, the majority of the microbial community at this site may not complete a single full cell cycle.

Differences observed between specific compounds indicate that specific taxonomic groups within a given soil exhibit turnover rates between 0.006 and 0.11 day^-1^, corresponding to generation times between 6.0 days and 1137.9 days (Supplementary Dataset S4). This range indicates highly heterogeneous growth rates across various constituents of the soil microbiome (Figure 1B). The majority of soil microorganisms in our study appear to be growing at extremely slow rates when compared to the maximum potential growth rates of many bacteria grown in culture (where generation times typically range from <1 to 100 h) (35). Although it is not surprising that the maximal growth of bacterial isolates in culture conditions do not represent *in situ* growth rates in soil, our finding that average, abundance-weighted generation times in soil microbial communities range from 20 – 64 days suggests that most soil microbes are oligotrophic, and studies of microbial growth *in vitro* may not necessarily be applicable to understanding microbial growth *in situ*.

Despite the slow growth rates inferred from our LH-SIP approach, these values should still be considered overestimates of ambient microbial growth rates. This is because the experimental conditions (water addition, mixing, and stable temperatures) necessarily provide more favorable conditions compared to *in situ* conditions. Despite this, both the community-level and compound-specific inferred growth rates of soil microbial communities prove to be exceedingly slow, further underlining the importance and ubiquity of slow-growing life in soil systems.

### Microbial biomass quantity does not predict growth

In all soils examined, we find no relationship between compound abundance and inferred growth rates (Figure 2). This indicates that rapidly growing taxa do not represent a large fraction of the soil microbial community at our field sites and, instead, most of the microbes found in bulk soil are relatively slow growing. Furthermore, on the time scale of our SIP incubation (0 - 7 days), growth of certain taxa did not clearly alter the bulk fatty acid profiles of the soils (Supplementary Figure 4), contrary to what one might expect if a minority of taxa were overgrowing the community. The uniformity of PLFA profiles throughout the incubation supports the utility of ^2^H_2_O as a tracer of *in situ* microbial growth, as there was no apparent modification of microbial growth with the addition of the tracer. In the three soils examined here, we observed large differences between the sites in the total quantity of PLFAs (ranging between 0 and 60 µg per g of soil), with the grassland and forest soils having smaller quantities of intact lipids (less microbial biomass) than the tundra soil. These differences in PLFA abundances are mirrored by differences in total organic matter for each of the soils (Supplementary Data 1). At the same time, growth rates were slower in the tundra soil (Figure 1B) and faster in the grassland and conifer forest soils. Although microbial biomass has been considered a proxy for the microbial productivity of a given soil (6, 7, 9, 10), here we find that total microbial biomass is unrelated to the rate at which this biomass is turning over. Our lipidomic dataset delineates two distinct soil microbiome profiles: a comparatively high turnover but lower biomass soil (typified by the conifer forest and grassland soils) and a relatively slower turnover but high biomass soil (the alpine tundra) (Figure 2). We suggest that soil microbiomes can be assessed along independent axes of biomass quantity and turnover, with the location of any given soil along these axes determined by biotic and abiotic conditions of the soil environment.

**Figure 2.**
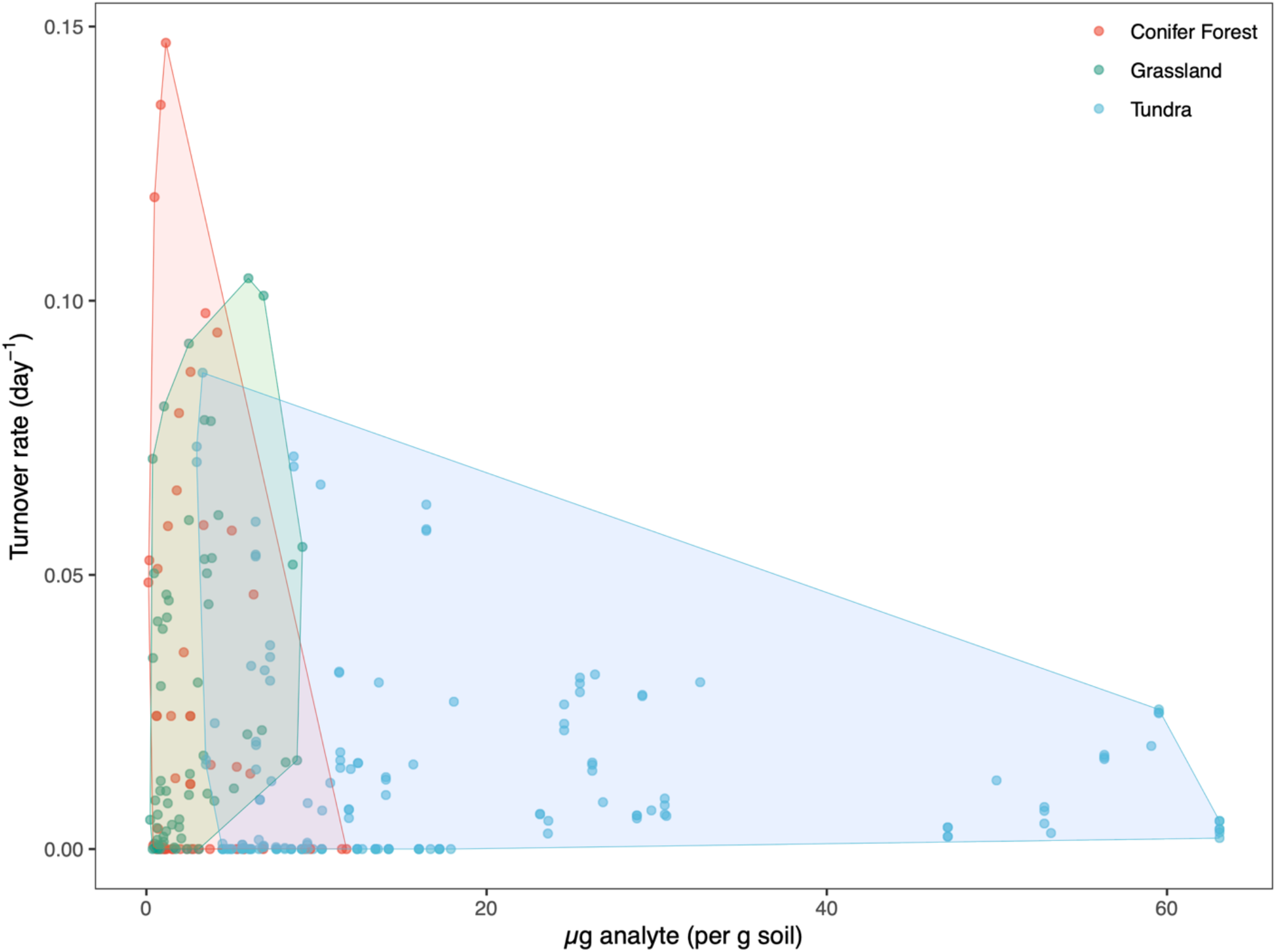
PLFA abundances, measured via GC-FID (µg per g soil), plotted against estimated turnover rates (day^-1^) in each of the three soils examined. These results highlight the lack of a relationship between compound-specific pool sizes and turnover rates in a given soil.

### Lipidomic data provides conservative taxonomic growth signals

Our 16S rRNA gene sequencing results (See *Materials and Methods*) show that the microbial communities in the soils examined were distinct but composed of bacterial taxa that are typically dominant in soils (Supplementary Figure 3): *Acidobacteria, Actinobacteria, Bacteroidetes, Chloroflexi, Proteobacteria, Planctomycetes*, and *Verrucomicrobia* (36). To examine whether any of these phyla could be distinguished with our lipidomic data, we mined the fatty acid profiles of 4959 taxa included in the Bacterial Diversity (BacDive) metadatabase (37). We observed that bacteria, at the phylum level, are broadly distinguishable based on their fatty acid profiles (Supplementary Figures 1 and 2), a finding supported by previous characterization of microbial PLFAs (38, 39). We also note that all major PLFAs we predicted to be present and representative of these phyla, based on our analysis of BacDive profiles, were indeed represented in our extracted lipid pools. We find that growth rate patterns by compound class are remarkably similar across all sample sites (Figure 1B). For example, terminally branched bacterial *iso*-15:0 and *iso*-17:0 saturated fatty acids consistently exhibited some of the fastest rates of turnover in each soil. These fatty acids are closely associated with the *Acidobacteria* and *Bacteroidetes* phyla (Supplementary Figure 1). Conversely, the 18:1 and 18:2 unsaturated fatty acids exhibited slower turnover rates. A large portion of the 18:1 and 18:2 unsaturated fatty acids likely represents the slower turnover of saprotrophic fungi (39–45) whose lifestyles may differ markedly from those of their bacterial neighbors.

We note that the LH-SIP method is limited in its taxonomic specificity due to the conserved nature of many groups of lipids (i.e., multiple bacterial taxa produce the same fatty acids) and the fact that the lipid profiles of some major soil bacterial taxa have not been well-characterized (31). For these reasons, our LH-SIP approach is not well-suited to determining community composition, but rather in quantifying the growth state of the community as a whole. Conservative inferences can be made regarding growth rates of bacteria and fungi at broad taxonomic levels based on the relative distributions of fatty acids, as demonstrated in previous studies (39). Thus, we provide the BacDive dataset (Supplementary Data 2) for the benefit of users interested in using our approach in soil and other systems. However, we emphasize that lipidomic datasets like these are best complemented by sequencing-based approaches to couple taxonomic information on communities to observed growth signatures.

### Comparing estimates of soil microbial growth

Published assessments of microbial growth rates in soil demonstrate disparate ranges that are highly dependent on methodology (12–20) (Figure 3). The LH-SIP results reported in this study yield assemblage-level growth rates between 0.01 – 0.03 day^-1^(generation times corresponding to 19.8 – 64.2 days) that are much slower than previously reported SIP methods with growth rates of 0.08 – 0.3 day^-1^(generation times of 2.4 - 10 days) (Figure 3, Supplementary Data 3, Supplementary Data 4). Much of this prior work has utilized SIP of isotopically labeled thymidine (TdR), leucine (Leu), or ^18^O-labeled water (H_2_^18^O) incorporation into nucleic acids or proteins (14–17, 46). A potential source of difference in results between these approaches is that DNA- and protein-based SIP methods are measuring faster growing organisms because DNA and protein are likely synthesized in greater abundance by organisms undergoing more rapid growth and division. TdR and Leu approaches could also stimulate growth through the provision of carbon and nitrogen in the tracer solution. Perhaps most importantly, the short incubation times typically used with these approaches (usually <48 h) likely lead to only those taxa with generation times shorter than the incubation times incorporating sufficient quantities of the isotopic tracer for detection. Microbial taxa with longer generation times may not progress through enough of their chromosomal replication cycle to incorporate sufficient tracer into their DNA to be distinguished by nucleic acid density fractionation. We propose that LH-SIP measurements of microbial growth provide a more conservative estimate of soil microbial growth that captures the apparent slow-growing majority of soil microbes. Because IRMS measurements can capture trace incorporation of ^2^H into the alkyl chains of membrane lipids, LH-SIP can capture extremely slow growth both at the compound-specific and bulk scale.

**Figure 3.**
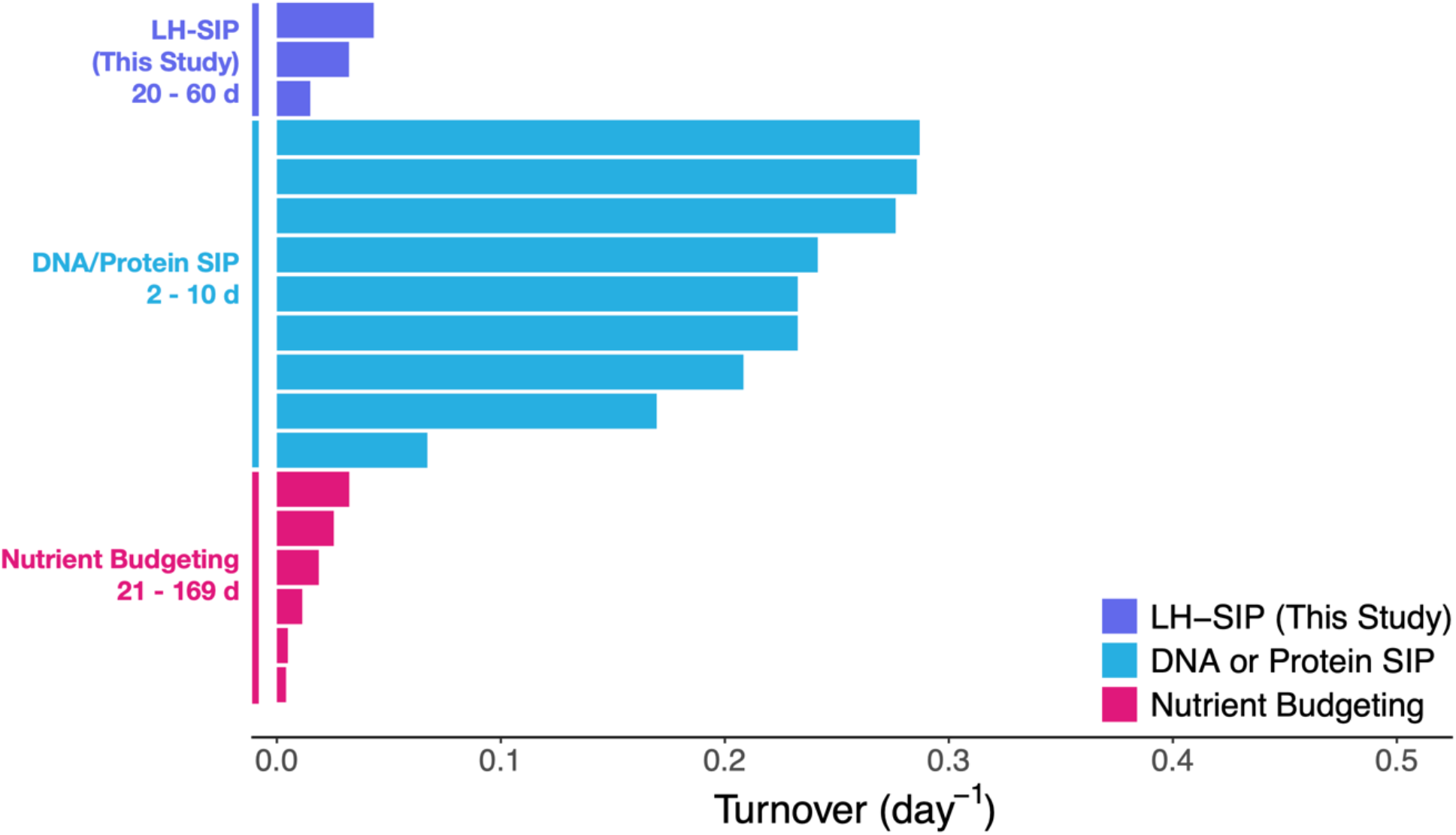
Published estimates of soil microbial growth rates (12–20) (collected as Supplementary Dataset 3), compared to those inferred by LH-SIP (this study). As expected, (see main text), we note that estimated growth rates with LH-SIP are much slower than those inferred by DNA/protein SIP techniques. Nutrient budgeting approaches, which typically measure the flux of carbon, nitrogen, or phosphorous through a soil to model microbial growth rate, generally lead to conservative estimates of microbial growth rates. Bar length corresponds to mean turnover/growth rate (day ^-1^) reported. Inferred generation times (days) are noted next to the label denoting the approach used.

### Future directions for LH-SIP

The growth rates inferred by LH-SIP of cells sharing lipid membrane constituents are aggregated. Single-cell methods including Raman spectroscopy or nanoSIMS (3, 24, 47) can elucidate cell-specific variation in growth rates while also measuring local mineralogical, elemental, or isotopic features. However, these methods can be resource intensive and often require separation of intact cells from environmental matrices, which can be problematic, especially in soil (48). A benefit of the LH-SIP approach is that it can provide community-level insights into anabolic growth activity that may be missed at the single-cell level. A two-pronged approach that couples bulk-scale LH-SIP measurements with single-cell metrics of growth heterogeneity could be powerful.

As noted above, LH-SIP is limited in its taxonomic specificity due to the conserved nature of many PLFA classes (39). Because LH-SIP yields compound-specific turnover rates, this method could be used in a highly targeted manner with systems containing less complex microbial communities or with strongly defined relationships between taxonomy and constituent lipid classes. Coupling LH-SIP with additional SIP (e.g., DNA or protein) or advanced lipidomic analyses has further potential to expand our understanding of the microbial physiology of slow growth in natural systems. For instance, future LH-SIP studies could take advantage of liquid chromatographic systems to elucidate relationships between intact polar lipid (IPL) head groups and associated growth rates. Alternatively, LH-SIP could be coupled with ^18^O-DNA qSIP to coregister highly sensitive measurements of LH-SIP with sequencing information of the more active fractions of the microbial community. Furthermore, the use of signature biomarkers for an organism or group of organisms in a given environment would allow for growth rate assignments with greater taxonomic specificity, potentially down to the species and strain.

### Relevance of the slow turnover of soil microorganisms

The *in vitro* cultivation and isolation of microorganisms from natural systems is notoriously difficult (13, 49, 50). There are numerous proposed reasons for this phenomenon, including the fact that media formulations are imperfect or select for certain taxa, but it is plausible that many “wild” microorganisms are not adapted for the rapid growth that is required for isolation and enrichment using standard cultivation approaches. Slow growth in soil may imply severe limitations on maximum growth rates even under ideal laboratory conditions, given that biochemical adaptations to slow growth may not be easily overcome in a lab environment. Many soil microorganisms in culture are observed to grow slowly, even in “ideal” conditions and, in fact, may be inhibited by high substrate concentrations (51–54). Our observed growth rates (Figure 1B) suggest that many soil microorganisms may be fundamentally difficult to cultivate due to time constraints on culturing experiments, as the amount of time for an organism to become visible on solid or in liquid media increases exponentially as doubling time increases (Supplementary Figure 6). We also emphasize that maximum potential growth rates measured *in vitro* likely do not reflect actual growth rates *in situ* as culture conditions may not adequately replicate the availability or paucity of carbon sources, electron donors/acceptors, and interactions with other organisms. Additionally, maximum potential growth rates (estimated via genomic analyses or culture-based experiments (35, 55)) are fundamentally different metrics than a direct measurement of growth in an environmental setting; a microbe capable of rapid growth under laboratory conditions will not necessarily exhibit this behavior under normal environmental conditions. This is highlighted by the observation that a wide array of microbial groups (including those with high maximum potential growth rates *in vitro*) persist in a dormant or nearly dormant state in soil (56).

## Conclusions

Here, we present evidence that slow microbial growth is widespread in soil systems. Across the soils analyzed, fatty acid abundances and growth rates were not correlated (Figure 2), indicating that the most abundant taxa are not necessarily the fastest growing. Soils with higher biomass can have slower growing communities, and vice versa. This result challenges the idea, often implicit in many studies documenting microbial biomass variation across soils, that higher biomass necessarily equates to higher soil productivity. Instead, our results suggest that soil microbiomes operate on a continuum of growth rate and biomass quantity, with the largest proportion of standing microbial biomass representing oligotrophic taxa adapted to slow growth. Growth rates presented here occur on the order of weeks to months, comparable to estimates generated by carbon- and nutrient-budgeting models (Figure 3). Our conclusion that slow-growing microorganisms appear to dominate the soil microbiome is in line with recent evidence that spatial variability in the composition of soil microbial communities typically exceeds the temporal variability observed at a given location (57, 58). Slow growth rates would be expected to attenuate short-term changes in overall microbial community composition, especially in soils with longer observed generation times. As microbial growth is a key regulator of a wide array of soil biogeochemical processes, our findings warrant additional studies that take advantage of the LH-SIP method described here to quantify variation in microbial growth rates across a broader array of soil types and conditions.

## Materials and Methods

### Soil sampling and incubation

Soils were collected from three locations in central Colorado: a conifer forest located at Gordon Gulch Critical Zone Observatory, Boulder County, CO (40.01, -105.46); a prairie grassland located at Marshall Mesa, Boulder County, CO (39.95, -105.22); and an alpine tundra located at Niwot Ridge, Niwot LTER (40.05, -105.58) near Ward, CO. The top 10 cm of soil were excavated with a surface-sterilized trowel, soils were sieved to 2 mm to remove rocks and plant material, and homogenized. Soils were stored in the dark at 4°C before incubations were started. For the SIP incubations, a 10 g subsample of each soil was weighed into a centrifuge tube and combined with 10 mL of filter-sterilized water with ∼5000 ppm (*δ*^2^H_VSMOW_ ≈ 31,000s ‰) ^2^H_2_O and incubated at 20°C for zero, three, or seven days. Isotopic composition of the incubation water was measured at the end of the time series experiment to account for the isotopic contributions of soil water (Supplemental Text). Samples were periodically shaken over the course of the incubation period to ensure uniform distribution of the tracer solution. At the end of the incubation period, excess incubation water was separated from the soil by centrifugation, decanted, and frozen for later isotopic analysis. Soil pellets were immediately flash-frozen by submerging in a dry ice ethanol bath and stored at -20°C until lipid extraction.

### Water hydrogen isotope analysis

The labeled incubation waters were analyzed for their H isotope composition (F_L_) after gravimetric dilution with water of known isotopic composition (1:1000 w/w) to get into the analytical range of available in-house standards previously calibrated to Vienna Standard Mean Ocean Water (VSMOW) and Standard Light Antarctic Precipitation (SLAP). 1µL of each sample was measured on a dual inlet Thermo Delta Plus XL isotope ratio mass spectrometer connected to an H-Device for water reduction by chromium powder at 850° (59). Measured isotope values in δ notation on the VSMOW-SLAP scale were converted to fractional abundances using the isotopic composition of VSMOW (R_VSMOW_ = ^2^H/^1^H = 0.00015576, (60)) and the relation F = R/(1+R) = (δ + 1) / (1/R_VSMOW_ + δ + 1) and corrected for the isotope dilution by mass-balance. The resulting isotopic composition of the tracer water was 5015 ppm ^2^H (31,357 ‰ vs. VSMOW). The isotopic composition of the labeled incubation water was diluted from this value after homogenization with the water in the soil depending on soil type and resulted in 4363 ± 251 ppm ^2^H for conifer forest, 4483 ± 49 ppm ^2^H for grassland, and 3492 ± 43 ppm ^2^H for tundra soils. The latter values were considered to be what cells encountered during tracer incubation (a combination of both the tracer and water present in the soil) and were used for all growth parameter calculations.

### Lipid extraction

Frozen soil pellets were lyophilized for 24 hours. Intact polar lipids were extracted from the dry pellets using a modified MTBE-based lipid extraction method (61, 62). In brief, 3.0 g of freeze-dried soil sample was added to a PTFE centrifuge tube. 3 mL of methanol was added to the sample and vortexed. 10 mL of MTBE was added to the sample and incubated at 1 h at room temperature while shaking. To induce phase separation, 2.5 mL of MS-Grade water was added and the mixture was centrifuged for 10 minutes at 1000G at room temperature. The organic phase was carefully extracted and transferred to an organic-clean glass vial. This process was repeated three times in total. Total lipid extract (TLE) was dried down under a stream of N_2_ gas and the sample was stored dry at -20 °C until solid phase chromatography. Prior to MTBE extraction, 100 μg of 23-phosphatidylcholine (23-PC) was added to all soils as an internal extraction standard. All glassware was furnaced at 450°C for 8 h prior to use. All Teflon vessels were solvent washed by sonication in a 9:1 mixture of DCM:MeOH for two sets of 30 minutes. Empty vessels were extracted alongside samples to monitor for contamination. No contamination was detected in the extraction blanks.

### Phospholipid separation and derivatization

Phospholipid extract (PLE) was purified from TLE using silica gel chromatography (62) to focus isotopic analyses on lipids derived from intact cells (free phospholipids outside of cellular membranes degrade relatively rapidly with half-life estimates of 39 hours at 15°C (63)). Combusted silica solid-phase extraction (SPE) columns containing 500 mg SiO_2_ were conditioned by the addition of 5mL acetone, then two additions of 5 mL dichloromethane (DCM). TLE was re-dissolved in 0.5 mL DCM and transferred to the SPE column. Neutral lipids and glycolipids were eluted by the addition of 5 mL of DCM or acetone, respectively, then dried down under a stream of N_2_ and stored dry at -20 °C. Phospholipids (PLE) were eluted by the addition of 5mL of methanol to the column. PLE was similarly dried under a stream of N_2_ and stored with an N_2_ -purged headspace at -20 °C.

The PLE was derivatized to fatty acid methyl esters (FAMEs) via base-catalyzed transesterification using methanolic base (64, 65). Transesterification was initiated by the addition of a mixture of 2 mL hexane and 1 mL 0.5 M NaOH in anhydrous methanol to dry PLE. The reaction mixture was allowed to proceed for 10 minutes at room temperature before being quenched by the addition of 140 μL of glacial (∼17 M) acetic acid and 1 mL water. The organic phase was extracted three times with 4 mL hexane and dried down under a stream of N_2_. A recovery standard of 10 μg 21-phosphatidylcholine (21:0 PC) was added to each PLE before derivatization to assess reaction yield. 10 μg of isobutyl palmitate (PAIBE) was added after derivatization to all samples as a quantification standard prior to analysis.

### FAME quantification and identification

A Thermo Scientific Trace 1310 Gas-Chromatograph equipped with a DB-5HT column (30 m x 0.250 mm, 0.10 µm) coupled to a flame-ionization detector (GC-FID) was used to quantify FAME concentrations (µg/g soil) and total amounts (µg extracted) based on peak area relative to the 23-PC extraction and PAIBE quantification standards, respectively. FAMEs were suspended in 100μL n-hexane and 1uL was injected using a split-splitless injector (SSL) run in splitless mode at 325°C; split flow was 12.0 mL per minute; splitless time was 0.80 min; purge flow was 5.00 mL/min; column flow rate was constant at 1.2mL/min. The GC ramped according to the following program: 80°C for 2 minutes, ramp at 20°C/min for 5 minutes (to 140°C), ramp at 5°C/min for 35 minutes (to 290°C). The FID was held at 350°C for the duration of the run. Major peaks were identified by retention time relative to a Bacterial Acid Methyl Ester (BAME) standard (Millipore-Sigma) and a 37 FAME standard (Supelco). Peak identities were confirmed using a Thermo Scientific Trace 1310 Gas-Chromatograph coupled to a single quadrupole mass spectrometer (ISQ) using identical injection and chromatography conditions with mass scans from 50-550 amu and a scan time of 0.2 seconds in positive ion mode (electron impact). FAMEs are referred to using the nomenclature z-x:y, where x is the total number of carbons in the fatty acid skeleton and y is the number of double bonds and their position (if known), while z-is a prefix describing additional structural features of the compound such as methylation and cyclization.

### FAME hydrogen isotope analysis

The isotopic composition of FAMEs was measured on a Thermo Scientific 253 Plus stable isotope ratio mass spectrometer coupled to a Trace 1310 GC via Isolink II pyrolysis/combustion interface (GC/P/IRMS). Chromatographic conditions were identical to those from the GC-FID and GC-MS stated above except for extension of the temperature program to baseline separate all major analytes (40°C hold for 2 minutes, 20°C/min to 120°C, then 2°C/min ramp to 240°C; 30°C/min ramp to 330°C, 4-minute hold) and injection via programmable temperature vaporization (PTV) inlet (ramped from 40°C to 400°C) to ensure quantitative transfer from the inlet to the column. Peaks were identified based on retention order and relative height based on coregistration with GC-FID and GC-MS chromatograms.

Measured isotope ratios were corrected for scale compression, linearity and memory effects using natural abundance and isotopically enriched fatty acid esters of known isotopic composition ranging from -231.2 ‰ to +3972 ‰ vs. VSMOW (see SI for details). Memory (compound-to-compound carryover) and linearity (peak-size) effects are important to correct for given the wide range of peak areas and isotopic values encountered in this study and the known impact of these effects on H isotope measurements (66, 67). Memory effect corrections for standard mixtures of natural abundance and enriched fatty acid esters lead to a >46% improvement in residual standard error (Fig. S7). Multivariable linear regression calibration including peak area terms with standards ranging in mass 2 areas from 1.74 Vs (∼25ng) to 69.67 Vs (∼975ng) lead to an overall RMSE of the calibration of 31.7‰ stemming from the substantial dynamic range of the isotope standards and believed to accurately reflect the elevated uncertainty that should be expected in the isotopically diverse samples. The conservative analytical standard errors ranged from 18.6 to 99.5 ‰ depending on peak area with larger error estimates for smaller peaks (see SI for details). The hydrogen sotope calibration was performed in R based using the packages *isoreader* (v 1.3.0 (68)) and *isoprocessor* (v 0.6.11) available at github.com/isoverse.

Calibrated isotope ratios measured via GC/P/IRMS were further corrected for H added during derivatization to FAMEs, as well as for analytical and replicate error as follows: First, the ^*2*^*F* of methanol used for base-catalyzed transesterification was measured by taking an aliquot of anhydrous methanol reagent (the exact stock used for transesterification) and derivatizing a phthalic acid with a known H isotopic composition (Arndt Schimmelmann, Indiana University) via acid catalysis. The resulting phthalic methyl ester was analyzed by GC/P/IRMS and a correction was applied to all values.

### Generation time calculations

Calculations of biosynthetic activity focus on the isotopic composition of FAMEs because the hydrocarbon skeleton of fatty acids consists only of C-H bonds that are nonexchangeable on biological time scales (69), unlike the readily exchangeable H bound to O, N, P and S in parts of lipid headgroups, proteins, and nucleic acids. The H tracer is thus stably incorporated into fatty acids tails during biological activity. The resulting isotopic enrichment of fatty acids in intact cellular lipids is described by the following equation (22):

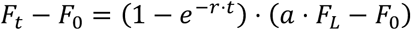

Where *r* is the specific biosynthesis rate [1/days]; *t* is the duration of tracer exposure [days]; *a* is the assimilation efficiency and fractionation of water hydrogen during lipid biosynthesis (see (22) and SI for details); and *F*_*0*_, *F*_*t*_, and *F*_*L*_ are the fractional abundances of ^2^H [ppm] in fatty acids before tracer incorporation, in fatty acids at time *t*, and in the isotopically labeled sample water. Solving this equation for *r* makes it possible to infer from the incubation time and isotopic measurements how quickly cellular fatty acids turn over. With lipid biosynthesis reflecting a combination of growth and repair, this provides an upper/lower bound for the specific growth rate *µ* and apparent generation time *T*_*G*_ Of the microbial producers of a given lipid:

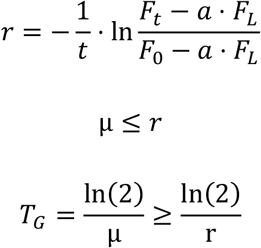

These calculations yield compound-specific generation time estimates that can be viewed by themselves or aggregated into assemblage-level estimates of community turnover. Uncertainty in each set of measurements was propagated through our calculations of µ by standard error propagation (see SI for details).

### 16S Ribosomal RNA Gene Sequencing

To characterize the microbial community composition of each soil type before SIP incubation, DNA was extracted from soil subsamples in triplicate and with negative control using the DNeasy PowerSoil DNA isolation kit (Qiagen, Germantown, MD, USA), according to the manufacturer’s instructions with one minor modification; samples were heated with Solution C1 for 10 minutes at 65 °C in a dry heat block prior to bead beating. Extracted DNA samples were amplified in duplicate using Platinum II Hot-Start PCR Master Mix (Thermo Fisher Scientific, Waltham, MA, USA) and the 16S rRNA gene primers 515F and 806R with Illumina sequencing adapters and unique 12-bp barcodes. The PCR program was 94°C for 2 minutes followed by 35 cycles of 94 °C (15 sec), 60 °C (15 sec), 68 °C (1 minute), and a final extension at 72 °C for 10 minutes. Amplification was verified via gel electrophoresis. Amplicons were cleaned and normalized with the SequalPrep Normalization Plate (Thermo Fisher Scientific) following manufacturer’s instructions and then pooled together. Sequencing was performed on an Illumina MiSeq using a v2 300 cycle kit with paired-end reads at the University of Colorado BioFrontiers Institute Next-Gen Sequencing Core Facility.

To prepare samples for analysis with the DADA2 (version 1.10.1) bioinformatic pipeline (70), reads were demultiplexed with adapters and primers removed using standard settings for cutadapt (version 1.8.1, Martin 2011). We used standard filtering parameters with slight modifications for 2×150 bp chemistry where forward reads were not trimmed and reverse reads were trimmed (truncLen) to 140 base pairs. In addition, we truncated reads at the first nucleotide with a quality score (truncQ) below 11 and a maximum allowed error rate (maxEE) of 1. These filtering parameters resulted in a mean of 95.7% of reads retained, and this was visually assessed with quality profiles for each sample. Reads were dereplicated, paired ends were merged, amplicon sequence variants (ASVs) were assigned, and chimeras were removed (98.23% of reads were not chimeric). Finally, taxonomy was assigned to each ASV against the SILVA (v132) reference database (71). We removed all chloroplast, mitochondria, and eukaryotic reads from the ASV table, which resulted in an average of 36029 reads per sample (range 27074 - 58528 reads), with the ASV table subsequently rarefied to 27000 reads per sample. Blank samples had far fewer reads than actual samples (mean of 193 reads compared to 35455 reads per sample), and the four genera detected in blanks (*Thermus, Geobacillus, Deinococcus*, and *Pseudomonas*) were not consistently detected and were below the 1% relative abundance threshold for inclusion in sample analyses. Taxonomic composition of the samples was compared across soil types (Supplementary figure 3).

### BacDive Database Analysis

To infer relationships between lipid profiles observed across our SIP incubations and high-level taxonomy, we queried the Bacterial Metadiversity Database (BacDive) (37) for all available fatty acid profiles using the BacDive API client implemented by the BacDiveR package (72). We generated a table of 4959 fatty acid profiles indexed by the taxonomy reported in the database and used principal component analysis to generate Supplementary Figure 2, grouped at the phylum level. We appended 24 manually curated fungal fatty acid profiles to this dataset, and this combined table is available as Supplementary Dataset 2. We looked at relationships between fatty acid profiles at the phylum level (Supplementary Figure 1) and conducted a principal component analysis of fatty acid composition and taxonomy (Supplementary Figure 2).

### Soil Geochemistry

Soils were analyzed at the Colorado State University Soil, Water, and Plant Testing Laboratory for routine determination of soil characteristics. In short, a KCl extract was used to quantify soil nitrate (73). An AB-DTPA extract was used to quantify soil P, Zn, Fe, Mn, Cu, and S (74). Organic matter percentages were calculated by determining the weight loss of samples after ignition. This data is available in Supplementary Dataset 1.

## Acknowledgements

We thank Julio Sepúlveda, Nadia Dildar, and Jonathan Raberg of the CU Boulder Organic Geochemistry Lab for valuable methodology input and Jessica Henley for assistance with 16S sequencing. We thank Jack Gugel, Toby Halamka, and Claire Karban for assistance with soil sampling. GC-FID, GC-MS, and GC-IRMS analyses were performed at the University of Colorado, Boulder Earth Systems Stable Isotope Lab (CUBES-SIL) core facility, RRID:SCR_019300. We thank the Feng Lab at Dartmouth College for isotopic determination of heavy water tracer solutions. This project was supported by a CU Boulder Grand Challenge Seed Grant to SHK and NF, a grant from the U.S. National Science Foundation (#2131837) to NF, and a research grant from the Army Research Office (#78484-LS) to SHK. TAC was supported by a National Science Foundation Graduate Research Fellowship and through the IQ Biology Program of the BioFrontiers Institute at the University of Colorado, Boulder. Logistical support for Niwot Ridge access was provided by the Niwot Ridge LTER program (NSF DEB – 1637686) with thanks to William D. Bowman for assistance with site access. SDJ was supported by the Canyonlands Research Center Graduate Research Scholars Program. The Canyonlands Research Station is supported by The Nature Conservancy.

