## Supplementary Information for "Hydrogen stable isotope probing of lipids demonstrates slow rates of microbial growth in soil"

### Supplementary Information Text

### Calibrating the compound specific H isotope data

The H isotope composition of fatty acid methyl esters (FAMEs) was measured by GC-P-IRMS and corrected for scale compression, linearity, and memory effects using natural abundance and isotopically enriched fatty acid esters of known isotopic composition including the F8 standard (containing 8 fatty acid methyl and ethyl esters from C14 FAME to C20 FAEE from -231.2 ‰ to -166.7 ‰ vs. VSMOW; A. Schimmelmann, Indiana University); custom mixtures with moderately enriched fatty acid esters (containing up to 4 natural abundance and 4 enriched methyl/ethyl/ propyl/butyl esters with C14 FAME at -231.2 ‰, C18 FAEE at -214.2 ‰, C20 FAEE at -195.5 ‰, C24 FAME at -179.3 ‰; and C16 FAEE at +275.6 ‰, C16 FAPE at +449.3 ‰, C20 FAPE at +191.9 ‰, C20 FABE at +1.5 ‰ vs. VSMOW; A. Schimmelmann, Indiana University); and custom mixtures with highly enriched fatty acid esters (containing natural abundance and 3 enriched methyl esters with C14 FAME at +3677 ‰, C16 FAME at +3828 ‰, C20 FAME at +3972 ‰ vs. VSMOW; calibrated in-house standard). Memory (compound-to-compound carryover) and linearity (peak-size) effects are particularly important to correct for given the wide range of peak areas and isotopic values encountered in the samples from this study and the known impact of these effects on H isotope measurements (1, 2). Memory effects were evaluated as detailed below using 340 peaks from mixtures of natural abundance and enriched standards run at different concentrations with the same chromatographic method as the samples (2°C/min ramp), as well as at a faster chromatographic ramp (5°C/min) for comparison. Linearity was corrected for as detailed below using the natural abundance and heavily enriched standards measured with the same chromatographic method as the samples (2°C/min ramp) at regular intervals and different concentrations with 849 calibration peaks in total and *m/z* 2 peak areas across all calibration peaks ranging from 1.74 Vs to 69.67 Vs.

### Derivations

The isotopic composition of an analyte peak ($R_{peak}$) is measured vs. the $H_2$ reference gas ($R_{H2}$) and constitutes the combination of the peak ($m_{peak}$) with true isotopic composition of the analyte ($R_{analyte}$) and the memory effects ($m_{mem}$) from all prior peaks ($\sum_{i<peak} m_i \cdot f(t_i)$) and their isotopic compositions ($R_{mem} = \sum_{i<peak} R_i \cdot m_i \cdot f(t_i)$) weighted by a time-dependent ($t_i$) memory decay function *f*. The resulting mass balance highlights how memory effects impact measured isotope signal in an area- (or amplitude-) dependent manner as they matter most for small peaks that closely follow large, isotopically different peaks ($m_{peak} < m_{mem}$; $R_{analyte} \neq R_{mem}$) and least for large peaks that follow small or isotopically similar peaks.

$$
\begin{aligned}
R_{peak} &= \frac{m_{peak}}{m_{peak} + m_{mem}} \cdot R_{analyte} + \frac{m_{mem}}{m_{peak} + m_{mem}} \cdot R_{mem} \\
&= \frac{m_{peak}}{m_{peak} + \sum_{i<peak} m_i \cdot f(t_i)} \cdot R_{analyte} + \frac{1}{m_{peak} + \sum_{i<peak} m_i \cdot f(t_i)} \cdot \sum_{i<peak} R_i \cdot m_i \cdot f(t_i) \\
\frac{R_{peak}}{R_{H2}} &= \frac{m_{peak}}{m_{peak} + \sum_{i<peak} m_i \cdot f(t_i)} \cdot \frac{R_{analyte}}{R_{H2}} + \frac{1}{m_{peak} + \sum_{i<peak} m_i \cdot f(t_i)} \cdot \sum_{i<peak} \frac{R_i}{R_{H2}} \cdot m_i \cdot f(t_i) \\
&= \frac{R_{analyte}}{R_{H2}} - \frac{\sum_{i<peak} m_i \cdot f(t_i)}{m_{peak} + \sum_{i<peak} m_i \cdot f(t_i)} \cdot \frac{R_{analyte}}{R_{H2}} \\
&\quad + \frac{1}{m_{peak} + \sum_{i<peak} m_i \cdot f(t_i)} \cdot \sum_{i<peak} \frac{R_i}{R_{H2}} \cdot m_i \cdot f(t_i)
\end{aligned}
$$

Taking into consideration that true isotopic compositions of standards and samples are known/sought vs.

VSMOW ($R_{VSMOW}$) and switching to $\delta$ notation (does not introduce any approximations as no terms are dropped) yields:

$$
\begin{aligned}
\frac{R_{peak}}{R_{H2}} &= \frac{R_{analyte}}{R_{VSMOW}} \cdot \frac{R_{VSMOW}}{R_{H2}} - \frac{\sum_{i<peak} m_i \cdot f(t_i)}{m_{peak} + \sum_{i<peak} m_i \cdot f(t_i)} \cdot \frac{R_{analyte}}{R_{VSMOW}} \cdot \frac{R_{VSMOW}}{R_{H2}} \\
&+ \frac{1}{m_{peak} + \sum_{i<peak} m_i \cdot f(t_i)} \cdot \sum_{i<peak} \frac{R_i}{R_{H2}} \cdot m_i \cdot f(t_i) \\
\delta_{peak/H2} &= \frac{R_{peak}}{R_{H2}} - 1 \\
&= \frac{R_{analyte}}{R_{VSMOW}} \cdot \frac{R_{VSMOW}}{R_{H2}} - \frac{\sum_{i<peak} m_i \cdot f(t_i)}{m_{peak} + \sum_{i<peak} m_i \cdot f(t_i)} \cdot \frac{R_{analyte}}{R_{VSMOW}} \cdot \frac{R_{VSMOW}}{R_{H2}} \\
&+ \frac{1}{m_{peak} + \sum_{i<peak} m_i \cdot f(t_i)} \cdot \sum_{i<peak} \frac{R_i}{R_{H2}} \cdot m_i \cdot f(t_i) - 1 \\
&= \left(\delta_{analyte/VSMOW} + 1\right) \cdot \frac{R_{VSMOW}}{R_{H2}} - \frac{\sum_{i<peak} m_i \cdot f(t_i)}{m_{peak} + \sum_{i<peak} m_i \cdot f(t_i)} \cdot \left(\delta_{analyte/VSMOW} + 1\right) \cdot \frac{R_{VSMO}}{R_{H2}} \\
&+ \frac{1}{m_{peak} + \sum_{i<peak} m_i \cdot f(t_i)} \cdot \sum_{i<peak} \left(\delta_{i/H2} + 1\right) \cdot m_i \cdot f(t_i) - 1 \\
&= \delta_{analyte/VSMOW} \cdot \frac{R_{VSMOW}}{R_{H2}} - \frac{\sum_{i<peak} m_i \cdot f(t_i)}{m_{peak} + \sum_{i<peak} m_i \cdot f(t_i)} \cdot \delta_{analyte/VSMOW} \cdot \frac{R_{VSMOW}}{R_{H2}} \\
&+ \frac{\sum_{i<peak} \delta_{i/H2} \cdot m_i \cdot f(t_i)}{m_{peak} + \sum_{i<peak} m_i \cdot f(t_i)} - \frac{\sum_{i<peak} m_i \cdot f(t_i)}{m_{peak} + \sum_{i<peak} m_i \cdot f(t_i)} \cdot \left(\frac{R_{VSMOW}}{R_{H2}} - 1\right) + \frac{R_{VSMOW}}{R_{H2}} - 1
\end{aligned}
$$

Introducing a transfer function $H(A_{peak})$ that relates the $1/m_{peak}$ area terms to the peak area $A_{peak}$ (rarely a linear transformation, more commonly takes forms such *log(A), 1/A,* $1/A^2$*, sqrt(A), etc.*) and rearranging including memory effect term $\delta_{mem/H2}$ provides the following multivariate linear regression:

$$
\begin{aligned}
\delta_{peak/H2} - \delta_{mem/H2} &= \frac{R_{VSMOW}}{R_{H2}} - 1 + \delta_{analyte/VSMOW} \cdot \frac{R_{VSMOW}}{R_{H2}} \\
&- H(A_{peak}) \cdot \left(\frac{R_{VSMOW}}{R_{H2}} - 1\right) - H(A_{peak}) \cdot \delta_{analyte/VSMOW} \cdot \frac{R_{VSMOW}}{R_{H2}} \\
&= \beta_0 + \beta_1 \cdot \delta_{analyte/VSMOW} + \beta_2 \cdot H(A_{peak}) + \beta_3 \cdot H(A_{peak}) \cdot \delta_{analyte/VSMOW}
\end{aligned}
$$

with:

$$
\begin{aligned}
\delta_{mem/H2} &= \frac{\sum_{i<peak} \delta_{i/H2} \cdot m_i \cdot f(t_i)}{m_{peak} + \sum_{i<peak} m_i \cdot f(t_i)} \\
H(A_{peak}) &= \frac{\sum_{i<peak} m_i \cdot f(t_i)}{m_{peak} + \sum_{i<peak} m_i \cdot f(t_i)}
\end{aligned}
$$

### Parameters

The best fit functional form and decay constant of the memory effects term $\delta_{mem/H2}$ was evaluated with data from mixtures of natural abundance and enriched standards run both with slow and fast chromatographic temperature ramps (2°C/min; 5°C/min) using numerical minimization of the residual standard deviation of the above multivariate linear regression with 340 standard peaks. The following functional forms for the memory decay $f(t_i)$ of preceding peaks' signals were evaluated and the exponential decay function was found to lead to the largest reduction in residual standard deviation (>46%) and most symmetric residuals for both 2°C/min and 5°C/min temperature ramps (see Fig. S7) with half lives (*ln 2 / k*) of 80.9 seconds and 26.1 seconds, respectively, reflecting the different time scales of peak elution for the different ramps.

$$\Delta_t = t_{\text{peak}} - t_i$$
$$f(t_i) = e^{-k\cdot\Delta t}$$
$$f(t_i) = 1 - \tanh k \cdot \Delta t$$
$$f(t_i) = \frac{1}{1 + k\cdot\Delta t}$$
$$f(t_i) = \frac{1}{(1 + k\cdot\Delta t)^2}$$

The raw data from all standard and sample analyte peaks (all run with 2°C/min chromatographic temperature ramps for improved peak resolution) was thus corrected for memory effects using the following equation:

$$\delta_{mem/H2} = \frac{\sum_{i<peak} \delta_{i/H2} \cdot A_i \cdot e^{-\ln 2\Delta t/80.9s}}{A_{peak} + \sum_{i<peak} A_i \cdot e^{-\ln 2\Delta t/80.9s}} = \frac{\sum_{i<peak} \delta_{i/H2} \cdot A_i \cdot 2^{-\Delta t/80.9s}}{A_{peak} + \sum_{i<peak} A_i \cdot 2^{-\Delta t/80.9s}}$$

To finally determine $\delta_{analyte/VSMOW}$ for all sample peaks, the following multivariate linear regression (derived above) was inverted after fitting to the 849 standard peaks run interspersed with the samples:

$$\delta_{peak/H2} - \delta_{mem/H2} = \beta_0 + \beta_1 \cdot \delta_{analyte/VSMOW} + \beta_2 \log A_{peak} + \beta_3 \cdot \log A_{peak} \cdot \delta_{analyte/VSMOW}$$

The overall RMSE of the calibration was relatively large at 31.7‰ from the substantial dynamic range of the isotope standards (-231.2 to +3972 ‰ and 1.74 Vs to 69.67 Vs areas) and we believe accurately reflects the elevated uncertainty that should be expected in the samples as well. The conservative analytical standard errors thus estimated by the inversion of this calibration uses binomial proportion confidence intervals (Wald intervals) and ranged from 18.6 to 99.5 ‰ depending on peak area (larger error estimates for smaller peaks).

### Correcting for the isotopic composition of the derivatizing agent

Fatty acid esters (e.g. phospholipids, glycolipids, and triglycerides) can be trans-esterified in the presence of a basic catalyst in anhydrous methanol to make fatty acid methyl esters (FAMEs). The reaction of anhydrous methanol (MeOH) with sodium hydroxide (NaOH) produces sodium methoxide (NaOH-MeOH) that is the trans-esterification agent. The isotopic composition of the fatty acid methyl esters produced from this process must be corrected for by taking into account the contribution of anhydrous methanol. The isotopic composition of anhydrous methanol is determined by derivatization to a phthalic acid methyl ester (PAME) by acid catalysis with acetyl chloride. The isotopic enrichment of PAME ($^2F_{PAME}$) is determined by isotope ratio mass spectrometry as described in *Materials and Methods*. The correction to the FAME measurement is therefore applied by mass balance. The measured isotopic fractional abundance of the FAME is the sum of the alkyl and methyl components weighted by their mass fraction.

$$^2F_{FAME} = {}^2F_{alk} \cdot x_{alk} + {}^2F_{Me} \cdot x_{me}$$

where $^2F_{alk}$ is the corrected isotopic composition of the alkyl chain, $^2F_{FAME}$ is the measured fractional abundance isotopic composition of the FAME, and x represents the mass fractions of the alkyl (alk) and the methyl groups (Me).

### Propagation of uncertainties

Measurements made in growth rate calculations carry with them varying levels of uncertainty. To account for this, we propagated error throughout our growth rate estimations as follows. These errors are notated as error bars in Figure 1B.

For the equation:

$$\boldsymbol{\mu = -\frac{1}{t} \cdot \ln\left(\frac{F_T - a \cdot F_L}{F_0 - a \cdot F_L}\right)}$$

We propagate the uncertainty in each variable by finding the partial derivatives of each measured variable:

$$\boldsymbol{\frac{\partial \mu}{\partial t} = -\frac{1}{t^2} \cdot \ln\left(\frac{a \cdot F_L - F_0}{a \cdot F_L - F_T}\right) \approx 0}$$

$$\boldsymbol{\frac{\partial \mu}{\partial F_0} = \frac{1}{t} \cdot \frac{1}{a \cdot F_L - F_0}}$$

$$\boldsymbol{\frac{\partial \mu}{\partial F_T} = -\frac{1}{t} \cdot \frac{1}{a \cdot F_L - F_T}}$$

$$\boldsymbol{\frac{\partial \mu}{\partial F_L} = \frac{1}{t} \cdot \left(\frac{a}{a \cdot F_L - F_0} - \frac{a}{a \cdot F_L - F_T}\right) = \frac{1}{t} \cdot \frac{a \cdot (F_0 - F_T)}{(a \cdot F_L - F_0) \cdot (a \cdot F_L - F_T)}}$$

$$\boldsymbol{\frac{\partial \mu}{\partial a} = -\frac{1}{t} \cdot \left(\frac{F_L}{a \cdot F_L - F_0} - \frac{F_L}{a \cdot F_L - F_T}\right) = \frac{1}{t} \cdot \frac{F_L \cdot (F_0 - F_T)}{(a \cdot F_L - F_0) \cdot (a \cdot F_L - F_T)}}$$

We apply these uncertainty terms by quadrature:

$$\boldsymbol{\sigma_\mu = \sqrt{\left(\frac{\partial \mu}{\partial t} \cdot \sigma_t\right)^2 + \left(\frac{\partial \mu}{\partial F_0} \cdot \sigma_{F_0}\right)^2 + \left(\frac{\partial \mu}{\partial F_T} \cdot \sigma_{F_T}\right)^2 + \left(\frac{\partial \mu}{\partial F_L} \cdot \sigma_{F_L}\right)^2 + \left(\frac{\partial \mu}{\partial a} \cdot \sigma_a\right)^2}}$$

We approximate the uncertainty in time (*t)* to be zero, given that the SIP experiments would have been rapidly quenched in a matter of minutes at the end of the incubation period. This simplifies our equation to:

$$\boldsymbol{\sigma_\mu = \sqrt{\left(\frac{\partial \mu}{\partial F_0} \cdot \sigma_{F_0}\right)^2 + \left(\frac{\partial \mu}{\partial F_T} \cdot \sigma_{F_T}\right)^2 + \left(\frac{\partial \mu}{\partial F_L} \cdot \sigma_{F_L}\right)^2 + \left(\frac{\partial \mu}{\partial a} \cdot \sigma_a\right)^2}}$$

Substituting in the previously defined partial derivatives yields:

$$\boldsymbol{\sigma_\mu = \frac{\sqrt{(aF_L - F_T)^2\sigma_{F_0}^2 + (aF_L - F_0)^2\sigma_{F_T}^2 + a^2 \cdot (F_0 - F_T)^2\sigma_{F_L}^2 + F_L^2 \cdot (F_0 - F_T)^2\sigma_a^2}}{t \cdot (aF_L - F_0) \cdot (aF_L - F_T)}}$$

The uncertainty in $F_L$, $\boldsymbol{\sigma_{F_L}}$ the isotopic enrichment of the label, can be altered by the water content of the soil. In our case, this resulted in minor but measurable dilutions of our label. We therefore estimated $\boldsymbol{\sigma_{F_L}}$ as the standard error of water isotopic measurements normalized to the mean isotopic measurement for each soil type.

Assimilation efficiency of the label, *a*, represents the fraction of lipid fatty acid hydrogen that is sourced from water (the label) as opposed to other sources. The uncertainty in assimilation efficiency of soil microorganisms, $\boldsymbol{\sigma_a}$, we estimate by constraining values of *a* reported in literature for heterotrophic organisms. We assume that the bulk of our soil community adheres to an assimilation efficiency coefficient representative of heterotrophy, typically reported to exist between 0.4 and 0.9. For $\boldsymbol{\sigma_a}$ we use the standard error of assimilation efficiencies between growth water and lipids reported in literature (Fig. S8).

The uncertainty in the isotopic enrichment of biomass at time $t = 0$, $\boldsymbol{\sigma}_{F_T}$ is approximated by the standard error in FAME internal standards 21:0 PC and 23:0 PC at time zero. Uncertainty in biomass at time $t$ $\boldsymbol{\sigma}_{F_T}$ is assessed as the standard error in isotopic measurements after applying $H_2$ calibrations and memory effect corrections as outlined above.

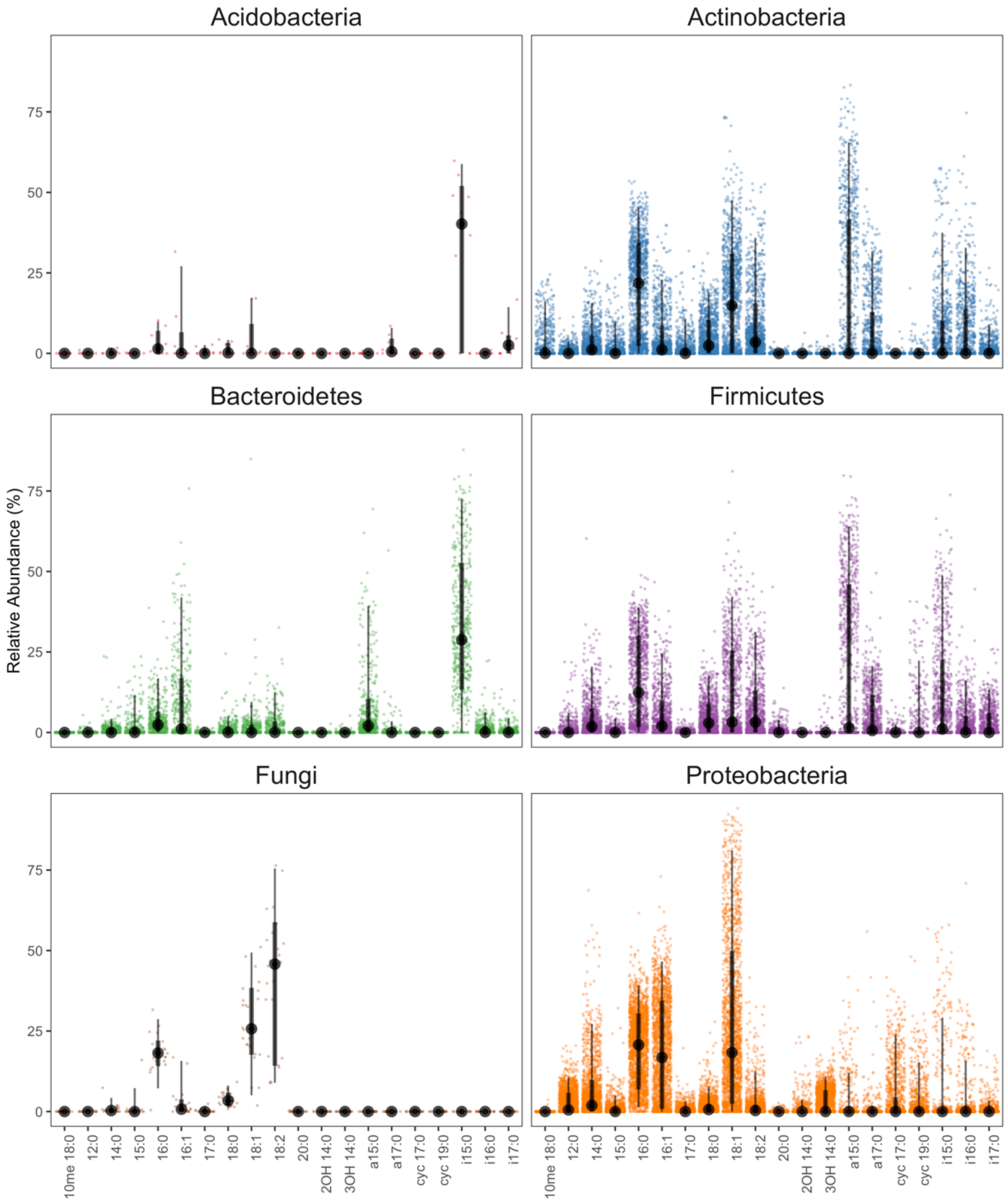


**Fig. S1.** Fatty acid profile data (n = 4959) mined from BacDive shows phylum-level taxonomic distinctions. Dots represent a single fatty acid measurement belonging to an organism of the indicated phylum. Large dots represent median values, thick bars represent the interquartile range, and thin bars represent 95% CI. Data used to generate this figure is available in Supplementary Dataset 2.

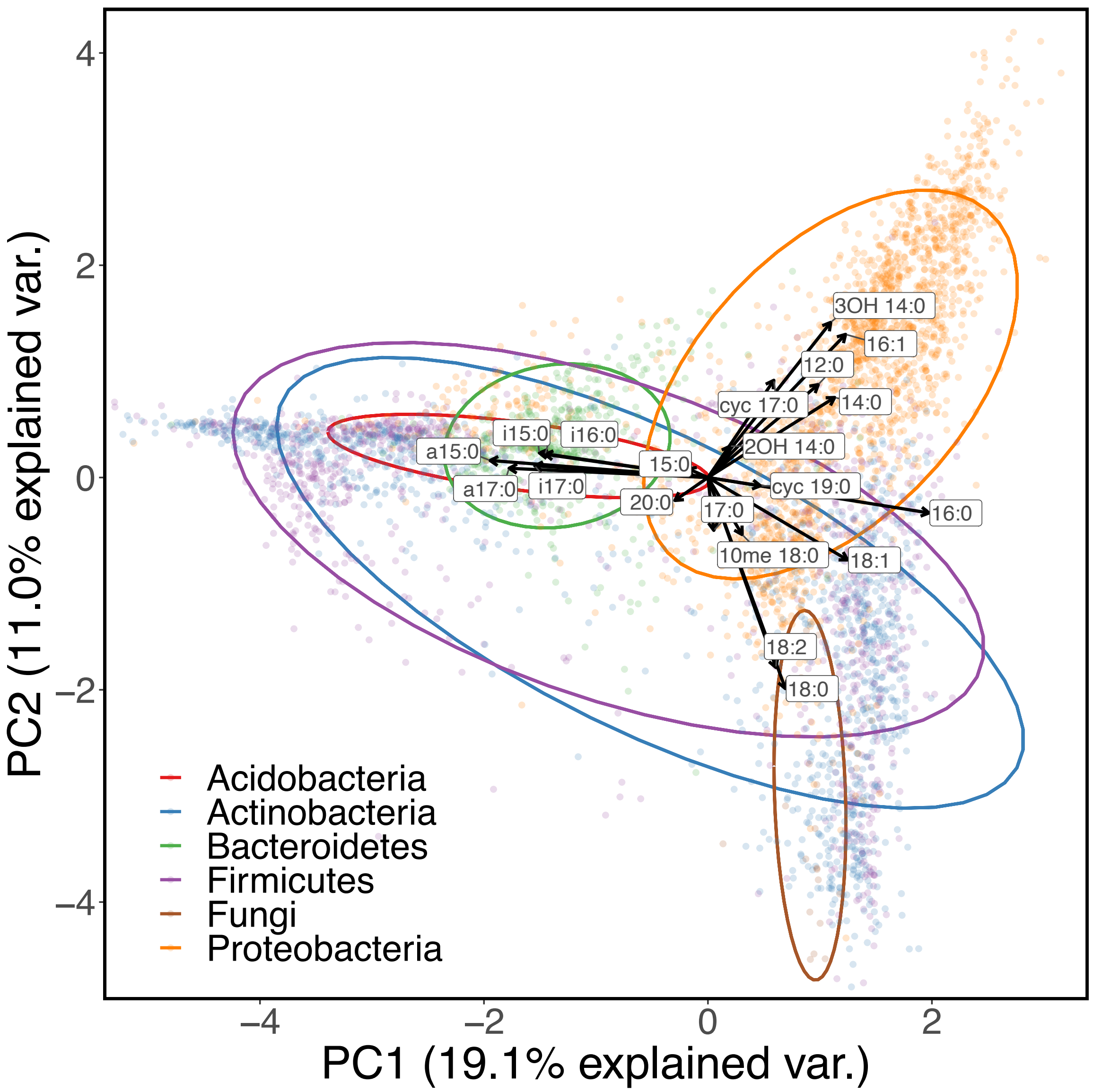


**Fig. S2.** Principal component analysis of fatty acid profile data (n = 4959) grouped at the phylum level mined from BacDive. Data used to generate this figure is available in Supplementary Dataset 2.

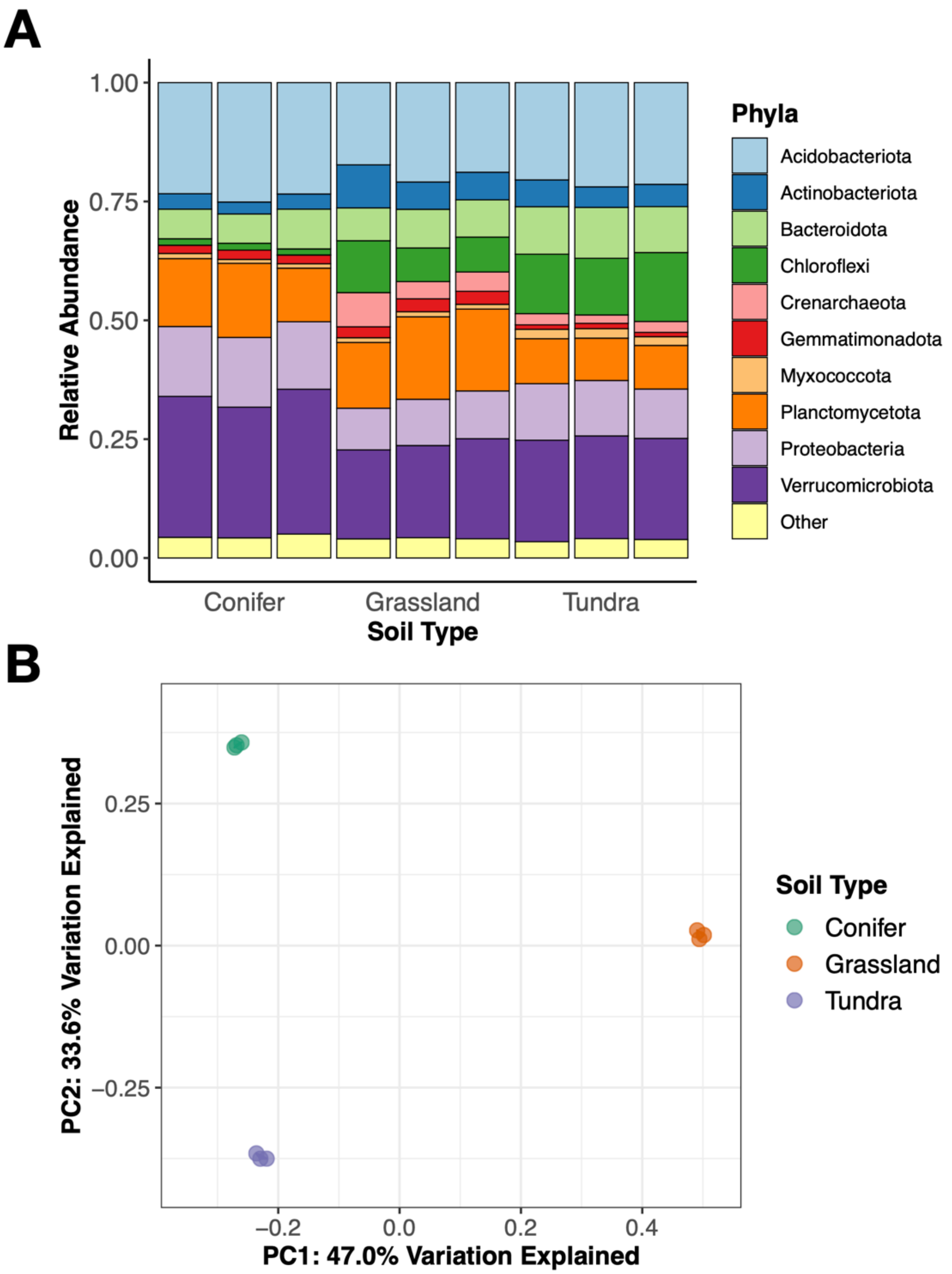


**Fig. S3.** 16S rRNA amplicon sequencing shows broadly similar but distinguishable soil microbial communities. Microbial phyla (A) are reported for each sample. Soils from each site were sequenced and are reported in triplicate. Principal component analysis (B) indicates that, while the major phyla between the soils are conserved, the soils examined harbor taxonomically distinct microbial communities.

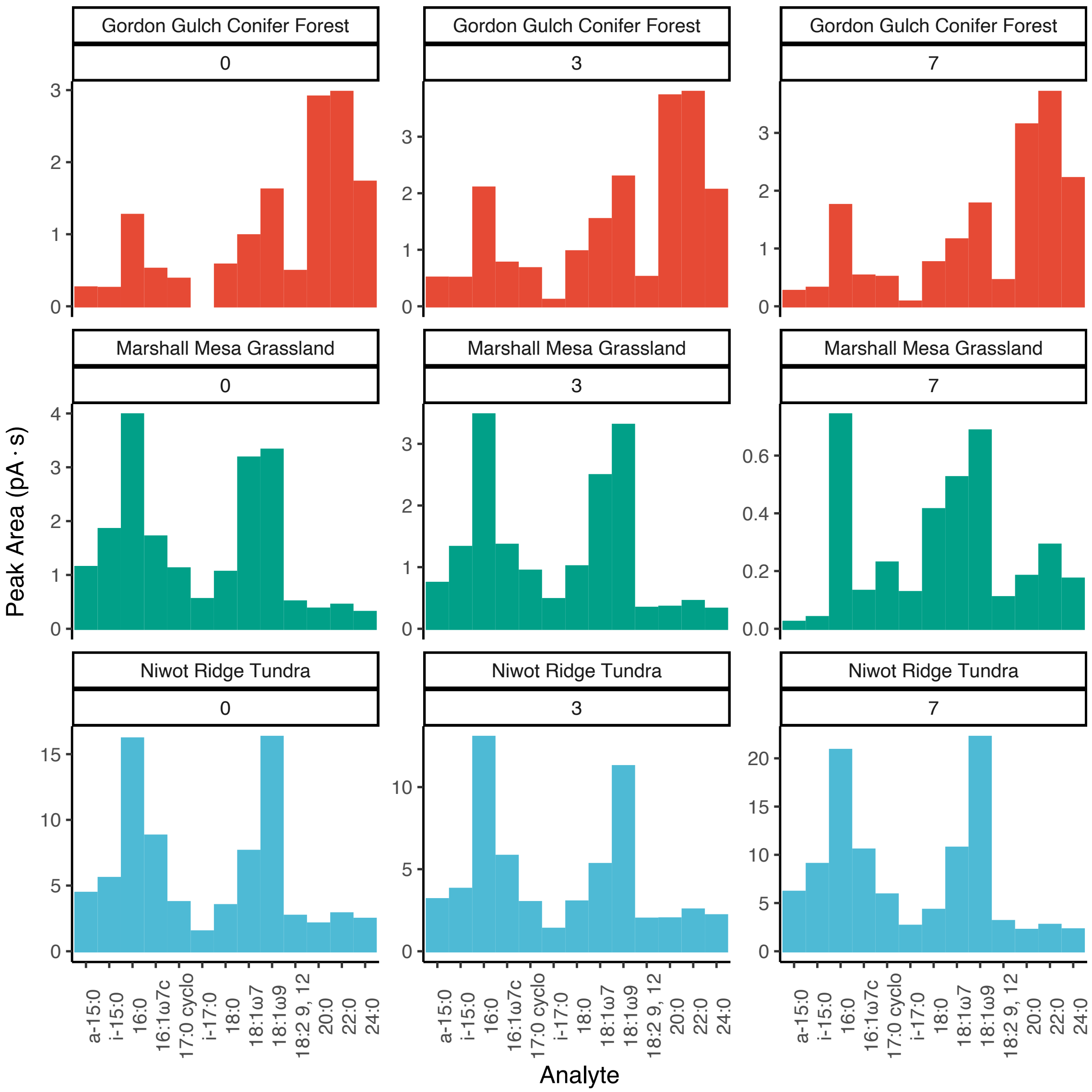


**Fig. S4.** Fatty acid profiles of soils included in this study, as measured by gas chromatograph flame ionization detector (GC-FID). Raw peak areas are reported in pA * second. Plot panels are numbered based on the day in the time series at which the soil was sampled.

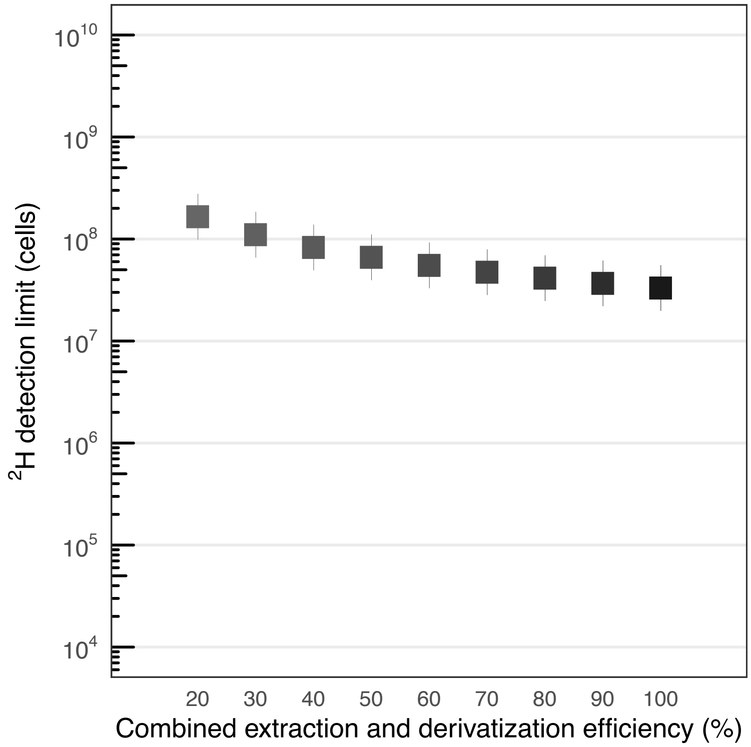


**Fig. S5.** Detection limit estimates of LH-SIP in units of minimum cells required for accurate $^{2}$H quantification via GC/P/IRMS. Detection limits are calculated for a variety of combined extraction and derivatization efficiencies (20 – 100%). The range of the bars represents a range of previously reported PLFA to cell conversion factors (3) in soil.

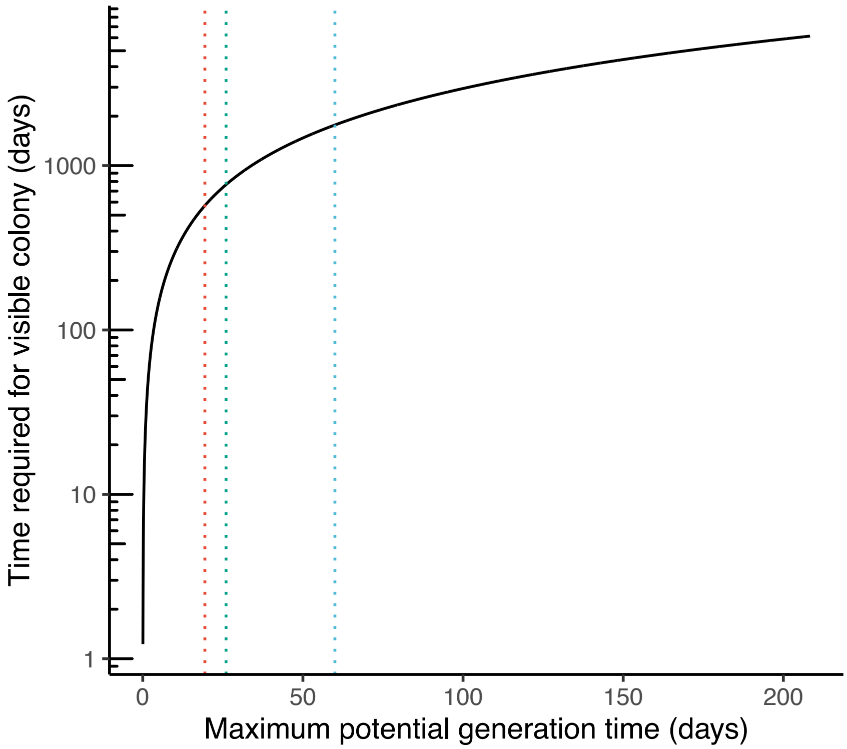


**Fig. S6.** Estimates of the time required to observe a visible colony (~1mm$^2$ ellipsoid) as a function of maximum potential generation time suggest that generation times observed in nature may preclude many taxa from culture-based techniques. Assumptions in this model include (i) cells do not produce biofilms or exudates, (ii) are continuously growing at an exponential rate, and (iii) are cultivated on an idealized growth medium (i.e. exhibit their maximum potential growth rate). Dotted vertical lines indicate the average apparent generation times estimated in the soils tested in this study: conifer forest, grassland, tundra (from left to right).

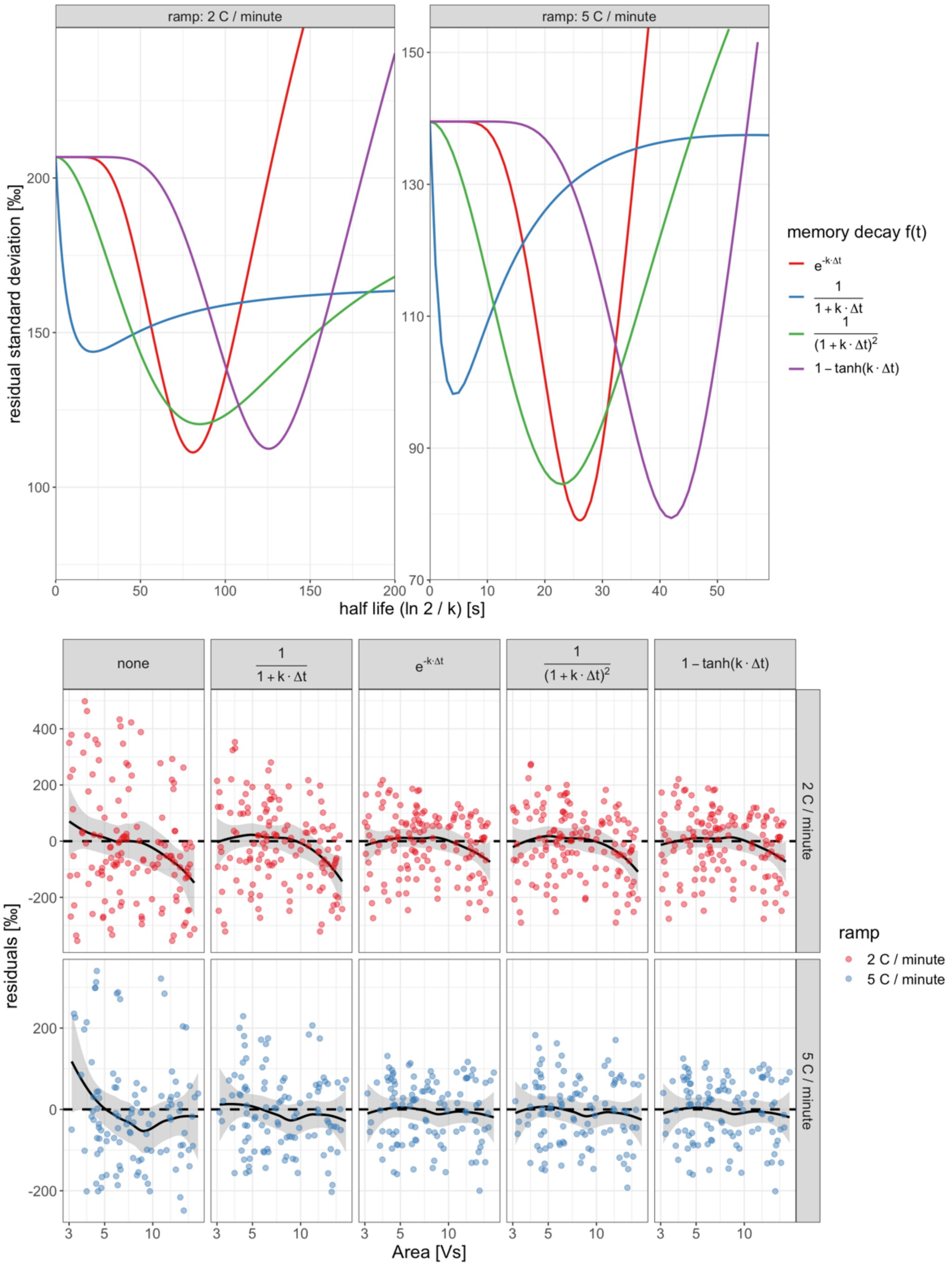


**Fig. S7**. **Top**: Improvements in residual standard deviation of multivariate linear regression $\delta_{peak/H2} - \delta_{mem/H2} = \beta_0 + \beta_1 \cdot \delta_{analyte/VSMOW} + \beta_2 \cdot H(A_{peak}) + \beta_3 \cdot H(A_{peak}) \cdot \delta_{analyte/VSMOW}$ with different functional forms for the temporal decay of the memory effect (signal carry over from preceding peaks) as a function of the decay constant *k* (1/s). *k* is displayed for clarity on the x-axis as the transformation *ln 2 / k* (in seconds), which is identical to "half-life" for the exponential decay form. Local minima indicate the best fit decay constants for each decay function. **Bottom**: residuals from the linear regression at the optimal decay constant for each functional form.

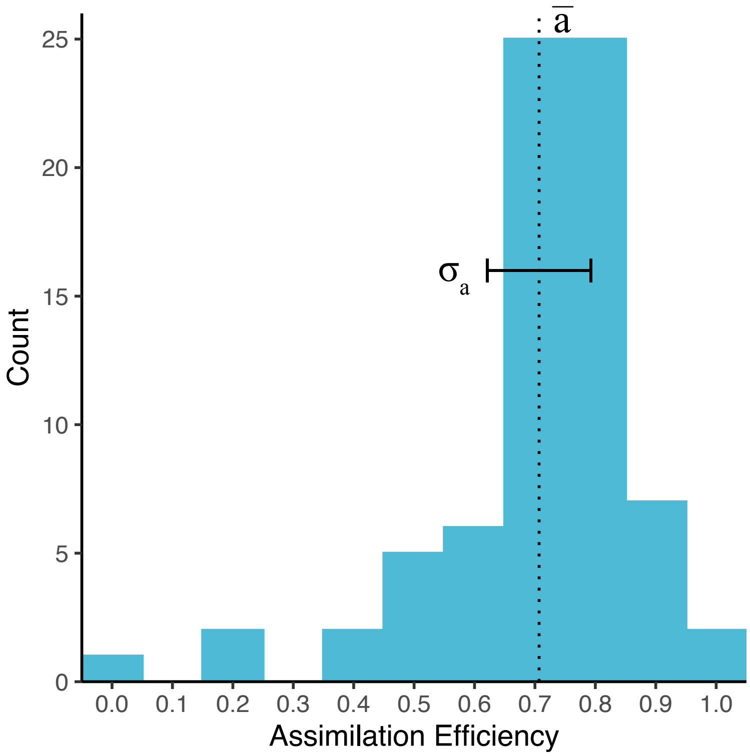


**Fig. S8.** Literature-reported assimilation efficiencies *(a)* between growth water and biomass lipid isotopic content cluster around 0.7. The mean *a* and $\sigma_a$ are the assimilation efficiency mean and standard deviation used in our growth rate calculations. The histogram values displayed are filtered to only include organisms utilizing heterotrophic metabolism.

### Supplementary Datasets

Dataset S1 (separate file). Edaphic information for each of the three soils included in this study as an excel workbook (.xlsx).

Dataset S2 (separate file). BacDive (4) fatty acid profiles aggregated at the phylum level as an excel workbook (.xlsx).

Dataset S3 (separate file). Literature-derived soil microbial growth rate values as an excel workbook (.xlsx).

Dataset S4 (separate file). Compound-specific hydrogen isotopic data and turnover values reported in this study, as an excel workbook (.xlsx).
